# *Trpv1^+^*sensory innervation of the salivary gland drives pain and supports saliva secretion

**DOI:** 10.64898/2026.08.06.743081

**Authors:** Deanna N. Cannizzaro, Juliana Amorim, Romy E. Wilson, Kayla M. Moehn, Akshayapriya Saravanan, Iva Vesela, Akash R. Gandhi, Isabelle M.A. Lombaert, Joshua J. Emrick

**Author notes:** Corresponding authors, Address: 1011 N. University Ave, Ann Arbor, MI 48109, Address: North Campus Research Center, Biointerfaces Institute, 2800 Plymouth Road, Ann Arbor, MI 48109. Equal contribution (Juliana Amorim, Romy E. Wilson).

## Abstract

Sensory neurons have been increasingly recognized as vital contributors to deep tissue function. However, how these specialized neurons contribute to salivary gland function remains largely undefined. Here, we uncover a role for trigeminal somatosensory afferents in salivary gland perception and function using *in situ*-based classification, *in vivo* calcium imaging, behavioral assays, and targeted ablation. Retrograde labeling from the submandibular gland complex revealed substantial direct innervation from trigeminal neurons. Further categorization confirmed that *Trpv1*^+^ sensory neurons provided dense innervation of the Wharton’s ducts. TRPV1 agonist ductal infusion directly activated gland complex-associated neurons in the trigeminal ganglia and evoked a robust pain phenotype. Targeted *Trpv1*^+^ ablation disrupted Wharton’s duct’s structure and dramatically reduced stimulated saliva volume. Our work provides the first evidence that *Trpv1*^+^ sensory neurons maintain salivary architecture and are necessary for stimulated saliva production, revealing a vital interoceptive role for direct trigeminal innervation in submandibular gland health.

## INTRODUCTION

Dynamic crosstalk between the somatosensory nervous system and peripheral target organs governs multiple aspects of tissue in health and disease. While peripheral sensory neurons in the dorsal root and trigeminal ganglia have canonically been defined in the context of external barrier tissues like the skin^1,2^, they are now known to also monitor deep internal tissue environments where they contribute to core functions therein through a process called interoception. For example, recent studies demonstrated that sensory neurons interact with the immune system to modulate inflammation and monitor tissue threats across diverse internal structures (e.g., lungs^3^, gastrointestinal tract, spleen, and lymph nodes^4^). These insights have fundamentally reshaped our understanding of somatosensation and interoception in the body, providing rationale for exploring additional deep tissues that are critical for maintaining oral health globally. While recent work has explored the trigeminal sensory systems that correspond to internal craniofacial tissues (e.g., hard palate^5^, tongue^5,6^, oral mucosa^5,7^, and teeth^8–12^), how defined sensory populations innervate and contribute to the saliva-secreting salivary glands remains poorly understood.

Saliva is an essential and versatile fluid that lubricates the oral cavity, protects against tooth demineralization, combats oral infection, and facilitates taste^13^. Three major paired salivary glands (i.e., parotid, submandibular, and sublingual) produce the bulk of saliva in mammals. Salivary glands comprise diverse cellular parenchyma, including acinar cells that produce and secrete saliva and epithelial ductal cells that carry and further modify saliva before its exit into the oral cavity^14^. Saliva secretion can be inhibited by ductal obstructions from salivary stones (i.e., sialolithiasis) and have been associated with acute orofacial pain^15,16^, hinting at a potential link with somatosensory afferents within one or more portions of the gland. Indeed, earlier rat studies supported the presence of trigeminal sensory innervation^17^ that may be nociceptive^18^. While these observations established a sensory presence and relevance in mammalian salivary glands, understanding of the identity and function of trigeminal subpopulations would represent a fundamental advance in exocrine gland biology and interoception.

Here, we uncover distinct trigeminal populations that target the submandibular glands’ complex (i.e., gland and associated Wharton’s duct), revealing a predominant *Trpv1^+^* sensory population. We followed up with comprehensive anatomical, physiological, and behavioral characterization of *Trpv1*^+^ afferents to define their contributions to sensation in the submandibular gland complex. Our work demonstrates that *Trpv1*^+^ fibers contribute to acute nociceptive pain and play a fundamental role in saliva secretion. Thus, our work defines a critical relationship between *Trpv1*^+^ trigeminal sensory innervation and the salivary gland complex and establishes a novel mechanism by which the sensory nervous system monitors and preserves internal craniofacial function.

## MATERIALS AND METHODS

### Animal assurance statement

All animal experiments were performed in accordance with protocols approved by the University of Michigan Institutional Animal Care and Use Committee following NIH guidelines. Experiments were performed with male and female mice. The number of mice used are indicated in the figure legends for each experiment. Mice were group-housed at room temperature with *ad libitum* access to standard lab mouse pellet food and water on a 12-hour (hr) light/12 hr dark cycle.

### Mouse lines

The mouse lines used in the present study included Ai95 [(RCL-GCaMP6f)-D (C57BL/6J)] (#028865, JAX), *Na_v_1.8*-Cre (#036564, JAX), C57BL/6J (#000664, JAX), Ai14 [(RCL-tdTomato)-D or LSL-tdTomato] (#007914, JAX), *Trpv1*-Cre (#017769, JAX), Ai140 [TIT2L-EGFP-ICL-tTA2] (#034100, JAX), and *Trpv1*-KO (#003770, JAX).

### Retrograde labeling of submandibular gland-innervating neurons using AAV6.2

Cannulation of Wharton’s duct for direct access to the submandibular gland and cell labeling has been previously described^19^. To fluorescently label neurons innervating the submandibular glands, adult (7-10 weeks of age) Ai14 (LSL-tdTomato reporter) mice were anesthetized via isoflurane administration (4% for induction and 2-3% for maintenance) using a SomnoSuite Low-Flow Anesthesia System (Kent Scientific) administered through a secured nose cone. Ophthalmic ointment (Fisher Scientific, Catalog #NC0490117) was applied to the eyes to prevent drying. The body temperature was maintained using a hand warmer (HotHands). Access to the Wharton’s duct was achieved with a custom-made aluminum breadboard platform (ThorLabs, MSB1015/M) where the maxillary incisors were hooked on a metal wire and the mandibular incisors were looped on a rope thread, which was gently pulled taut to hold the mouth agape. PE10 polyethylene tubing (Cat# 50-195-5495, Braintree Scientific) was heated and pulled to a compatible diameter for individual ducts (∼100 µm). The tubing was then manually guided with forceps (Item No. 11251-35, style #5/45, Fine Science Tools) for 3-5 mm into the submandibular gland’s Wharton’s duct, which is located at the center of the lingual papillae beneath the tongue. Tissue adhesive (Cat #1469SB, Vetbond) was applied around the cannula with an absorbent paper point (Meta, XX-fine) to keep the cannula in place. A 0.5 mg/mL of atropine sulfate salt monohydrate (Product #A0257) was intramuscularly injected at 1 µL/g body weight into each mouse and then waited at least 10 minutes to minimize saliva secretions that might disrupt the uptake of the infusions. The free end of the cannula tubing was then taped down to the breadboard for stability before inserting a custom-made Hamilton syringe (30-gauge needle, Model #705). Manual infusions included 50 µL of 1:1 sterile saline:ssAAV-6(F129L)/2-*hSyn1*-iCre-WPRE-SV40p(A) (Physical titer: 5.2×10E12 vg/mL, Catalog #223-6(F129L)/2, University of Zurich Viral Vector Facility VVF, Zurich, CH). Post-infusion, the syringe was left in the cannula for 10 min to prevent backflow before removing the cannula and placing the mouse back in its cage. Mice recovered for at least 3 weeks to ensure adequate viral-induced expression prior to euthanasia and tissue dissections.

### Pharmacologic TRPV1-mediated ablation using resinaferatoxin

Pharmacologic ablation of cells expressing TRPV1 in the submandibular gland was achieved via ductal delivery of resiniferatoxin (RTX) (#HY-N2333, MedChemExpress). RTX was prepared as described previously^20^. In short, a glass vial of RTX (1 mg) was solubilized in 150 μL ice-cold 100% ethanol, diluted with 50 μL ddH2O supplemented with ascorbic acid to make a final RTX stock concentration of 5 μg/μL RTX and 2 mM ascorbic acid. Stock RTX solution was further diluted to a working concentration of 100 ng/μL in Milli-Q water (0.25% Tween-80, 2 mM ascorbic acid, 0.9% NaCl). Vehicle was also used to dilute RTX (100 ng/μL) to achieve lower working concentrations. Wharton’s ducts were infused bilaterally in each mouse with a 50 ng dose (20 µL of a 2.5 ng/µL solution into each duct). Saliva collection and immunostaining were completed 24 hours post-RTX infusion, as described below.

### AAV6.2-*hSyn*-DTA-mediated ablation of *Trpv1^+^* neurons

Neurotoxic-based ablation of *Trpv1*^+^ neurons innervating the submandibular gland was achieved via infusion of ssAAV-6(F129L)/2-*hSyn1*-chl-dlox-DTA(rev)-dlox-WPRE-bGHp(A) (Physical titer: 3.6×10E12 vg/mL, Catalog #1014-6(F129L), University of Zurich Viral Vector Facility VVF, Zurich, CH). Wharton’s ducts were infused bilaterally in each mouse (20 µL of 1:1 sterile saline:AAV into each duct). Mice were allowed to recover for at least 3 weeks to ensure adequate viral-induced expression prior to saliva and histology evaluation.

### *In situ* hybridization (ISH) of trigeminal ganglia (TG) in tissue sections

Mice were euthanized and TGs were freshly dissected, flash frozen on dry ice, then embedded in OCT for cryosectioning (Leica CM1950) at a thickness of 20 μm onto Superfrost Plus Slides (Fisher #12-550-15). Slides were then used immediately or stored in the-80°C until use. In situ hybridization (ISH) was performed via an adaptation of hybridization chain reaction (HCR) version 3^21,22^.

Buffers, hairpin amplifiers, and probes against transcripts for mouse genes *S100b (PRA454), Calca (PRB722), Trpv1 (PRB721), Mrgprd (PRD964), Fxyd2 (PRB 40), Nppb (PRF446), Trpm8 (PRB982), Tmem233 (PRB735), Trpa1 (PRF447), and TdTomato (PRF678)* were purchased from Molecular Instruments. Slides were fixed in 4% PFA/1x PBS for 15 min on ice, PFA was chelated, then washed 3 times with 1x PBS before undergoing dehydration through an ethanol series (50%, 70%, 100%, fresh 100%; 5 min each) then rinsed 3 times in 2x saline sodium citrate (SSC). Next, slides were prehybridized with hybridization buffer using a parafilm at 37°C for 10 min in a humidified hybridization oven. This was followed by incubation with a working hybridization solution (prepared by adding 4 probes with adapters B1-B6 at a concentration of 4 nM per probe to pre-warmed hybridization buffer at 37°C), and incubated for 2-3 days in a humidified hybridization oven at 37°C. Following hybridization, slides were washed in 100% wash buffer for 5 min at 37°C, then underwent a wash solution series (75% wash buffer in 5x SSC containing 0.1% Tween 20 (SSCT), 50% wash buffer in 5x SSCT, 25% wash buffer in 5x SSCT, 100% 5x SSCT for 10 min at 37°C) with a final wash in 100% 5x SSCT for 5 min at room temperature (RT). Next, slides were pre-incubated in amplification buffer for 30 min followed by overnight incubation in a working amplification solution at RT in a humidified chamber protected from light. The working amplification solution consisting of amplification buffer containing amplifier hairpins for adapters B1-B6 conjugated to fluorophores was freshly prepared according to manufacturer recommendations, ensuring the selected associated fluorophores (Alexa 488, Alexa 561, Alexa 647, Alexa 750) had no overlap with any endogenous fluorescence present in the sample for each experiment. Finally, slides were washed 2 times in 5x SSCT for 15 min with gentle agitation, then mounted in Imaging Buffer (3 U/mL pyranose oxidase, 0.8% D-glucose, 2x SSC, 10 mM Tric HCl pH 7.4, 400 U RNAse inhibitor), coverslipped, and sealed for imaging using Cytobond (Scigene). The slides were imaged using confocal microscopy (Olympus FV3000, Evident Scientific, Inc.).

After imaging, sections were rinsed in 2x SSC, and DNA probes and amplifiers were removed by incubation in 250 U/mL RNase-free DNase (Roche Diagnostics) for 90 min at room temperature. Sections were washed 6 times (5 min in 2x SSC) and pre-hybridized for the next round of hybridization with the next 4 probes. This procedure was repeated for a total of 3 rounds of hybridization as previously described^12,21^. Automatic transcriptomic class assignment was carried out as described previously^21^.

### Immunohistochemistry of submandibular glands and Wharton’s ducts

For immunofluorescent staining, mouse submandibular glands and ducts were processed using distinct protocols. For submandibular tissue, glands were harvested and fixed in 4% paraformaldehyde (PFA) overnight at 4 °C, washed in 1x PBS, and cryoprotected sequentially in 15% and 30% sucrose. Samples were then incubated overnight in a 1:1 mixture of 30% sucrose and OCT compound, embedded in OCT on dry ice, and cryosectioned at 10 µm thickness. For duct analysis, a whole-mount protocol was used. Entire tissues were placed on filters (Nucleopore Track-Etch Membrane, Whatman #110405) in 1x PBS, equilibrated for 15 min at 37 °C, fixed in 4% PFA for 20 min at room temperature, and washed in 1x PBS. Indirect immunofluorescence was performed on both gland sections and duct whole mounts. Tissues were permeabilized with 0.1% Triton X-100 for 8 min and washed three times in 0.1% Tween-20. Samples were blocked for 2h at room temperature in blocking solution containing 10% donkey serum, 1% BSA, and 1.8% MOM IgG Blocking Reagent diluted in 0.1% Tween-20. Primary antibodies were applied overnight at 4 °C (Table 1). After washing, samples were incubated with Cy2-, Cy3-, or Cy5-conjugated secondary antibodies (Jackson Laboratories) and DAPI for 40 min at room temperature. Following final washes, gland sections were mounted in Fluorogel and coverslipped. Duct whole mounts were transferred to slides using secure-seal imaging spacers (Grace Bio-Labs #654004), mounted in Fluorogel, and coverslipped. Coverslip edges were sealed with nail polish. Imaging was performed using a Nikon A1R confocal microscope with z-stack acquisition at a step size of 0.957 µm to 5.75 µm for glands and 0.8 µm for ducts.

**Table 1.**
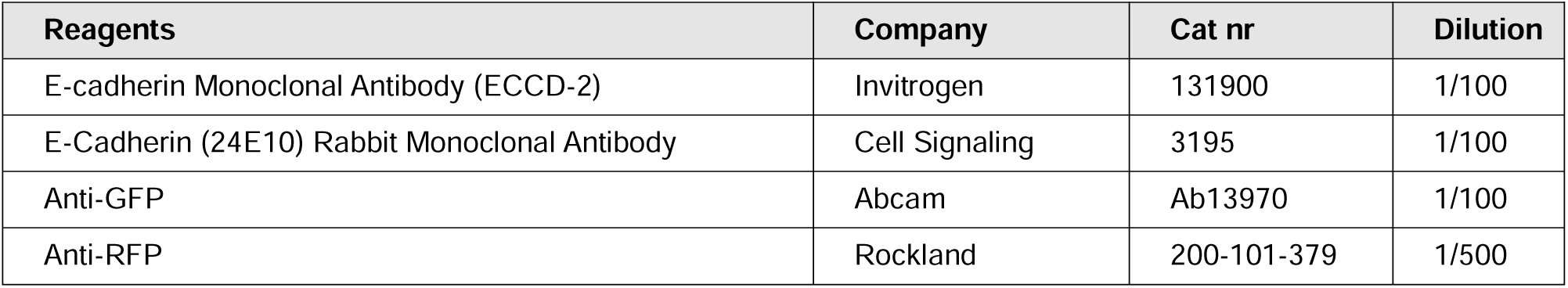
Antibodies used.

Gland and duct quantification was performed using QuPath software (v0.6.0). For the glands, a threshold was created to identify DAPI staining and define the number of cells within a region of interest (ROI). From the identified cells, both GFP^+^ and tdTomato^+^ cells were counted and the percentage of positive cells relative to total cells was calculated. For the innervation of Wharton’s ducts, a classifier was created using the threshold tool according to GFP staining. Positive nerve signal and background were annotated as distinct classes, and the classifier performance was iteratively refined using the live prediction tool to optimize signal discrimination. Once finalized, the classifier was saved and applied identically to all images to ensure consistency across samples. Ductal ROIs were defined based on DAPI staining to delineate the ductal area. Within each ROI, the trained classifier was applied to quantify positive green pixels within the ductal area. The percentage of innervation was calculated as the proportion of positive area relative to the total ductal area. Pixel detections were generated accordingly, and a network feature was created to extract quantitative parameters of innervation. Network density was numerically classified using the U-M generative AI, GPT 5.5. The metric considered a score ranging from 0 to 3, where 0 represented very low/near absent and 3 very high.

For Hematoxylin and Eosin (H&E) staining, glands were fixed in 4% PFA overnight, washed with 1x PBS (3 times), and preserved in 70% ethanol until processing and further slide scanning.

### *In vivo* epifluorescence Ca^2+^ imaging of the trigeminal ganglion

*In vivo* Ca^2+^ imaging of trigeminal neurons was performed in anesthetized mice using previously described methods^11,22^. Mice were anesthetized via isoflurane administration (4% for induction and 2-3% for maintenance) using a SomnoSuite Low-Flow Anesthesia System (Kent Scientific) administered through a secured nose cone. Ophthalmic ointment (Fisher Scientific Catalog #NC0490117) was applied to the eyes to prevent drying. Body temperature was maintained using a hand warmer (Hothands). Mice had one Wharton’s duct cannulated, as previously described, while being head-fixed using a custom stereotaxic apparatus that stabilized the skull while enabling access to the oral cavity. Optical access of the trigeminal ganglion surface was achieved via bilateral hemispherectomy. Following trigeminal ganglion exposure and establishment of hemostasis, Ca^2+^ imaging was performed within 30 min of trigeminal ganglion exposure using a Thorlabs custom-built epifluorescence microscope with a 4x, 0.16 NA, Olympus objective. A white light LED Controller (Prizmatix) was used to image GCaMP6f (480 nm) and RFP (561 nm). Fluorescent light was filtered using a GFP and RFP filter set (Thorlabs). Image acquisition was performed using a pco.panda 4.2 bi scientific CMOS camera at 5 Hz. For all experiments, a piezoelectric actuator (NanoScan NPC-D-6111, Prior) was used to achieve highly precise, stepwise, vertical linear movement of the Olympus objective in order to capture a deeper field of view (i.e., 300 μm) of the surface of the ganglion. A total volume of 10 µL of either RTX or vehicle was infused with a controlled, stereotaxic injector (Stoelting, QSI Dual Microliter Syringe Pump, Catalog #53325) at a rate of 3 µL/min. A custom-made Hamilton syringe 30-gauge needle (Model #705, point style 4, 10 mm length, 45-degree angle) was inserted into the cannula and placed into the injector. Videos were captured with 2,000 frames, including baseline from 0-500 frames and liquid infusion lasting from 3.3 min (approximately 550-1,550 frames).

### Calcium Imaging Analysis

Calcium imaging analysis occurred, as previously described^11^.

### Regions of interest (ROIs) selection

Regions of interest (ROIs) were manually drawn using the freehand tool in ImageJ around neurons that fluoresced during the applied infusion.

### Fluorescence dynamics

As previously published, a MATLAB script^23^ was adopted to calculate GCaMP6f fluorescence (ΔF/F) and correct for fluctuations in background signals for the manually drawn ROIs. A neuropil region local to each somatic ROI was created to enable spatial averaging of the Ca^2+^ response, then subtraction from the somatic average occurred. This process was repeated for every frame as previously described^11^. Analyzed data were used to generate heatmaps using MATLAB.

### a) Quantification analysis

GCaMP6f fluorescence traces (ΔF/F) were analyzed using a custom MATLAB script (a-d) (https://doi.org/10.24433/CO.5711439.v1). For each recording, the first 500 frames were excluded to remove baseline drift artifacts prior to analysis. Neurons were classified as active (i.e., responders) if the fluorescence signal reached or exceeded a threshold of ΔF/F ≥ 5 for a minimum of three consecutive frames. The frame at which each neuron first met this criterion was defined as the onset frame.

### b) Peak ΔF/F and AUC

To quantify response magnitude, the peak ΔF/F value was identified for each neuron post-onset window. The duration of calcium response was estimated by calculating the area under the curve (AUC) per minute for each neuron’s post-onset fluorescence trace. Median values are displayed for each condition.

### c) Linear regression

Linear regression was performed on peak ΔF/F versus AUC to assess the relationship between response magnitude and duration of calcium activity across vehicle and RTX conditions.

### d) Calcium traces

To visualize the range of individual neuron dynamics per condition, raw fluorescence traces were extracted from a window spanning 10 seconds before and 60 seconds after each neuron’s individual onset frame.

### Behavioral analysis of TRPV1 activation in the submandibular glands via RTX

C57BL/6J were habituated individually in a Plexiglas acrylic chamber (3.78″ inner diameter x 6″ height) equipped with a custom built controllable rotating stage inside a sound attenuating box (Item # ENV-017M-27, Med Associates Inc.). An infrared light (Item #A14, Tendelux) provided illumination from the ceiling. A monochrome camera (Blackfly S USB3 Model: BFS-U3-13Y3M-C, 1280P, 30 fps) equipped with a fixed focal length lens (#33-307, 8mm UC Series, Edmund Optics) was positioned horizontally facing the stage to capture mouse behaviors and facial changes. Habituation to the recording chamber took place for at least three sessions, 30 min each. Following habituation, each animal underwent a total of 2 experimental paradigms on separate days. The same cohort of mice was assayed for both conditions (vehicle then RTX, see below). Cannulation procedure was followed, as described above, without the use of atropine. Saline drops were administered to the eyes for lubrication instead of ointment for video tracking. Mice were first evaluated with 10 µL of vehicle (0.25% Tween-80, 2 mM ascorbic acid, 0.9% NaCl) infused into one Wharton’s duct under isoflurane. To assess behavioral responses to peripheral TRPV1 activation, 10 µL of RTX was infused into the same Wharton’s duct (50 ng dose as described above) 2-3 days after vehicle infusion. Following infusion, mice stayed under anesthesia for 2 min to prevent backflow of liquid. Cannulas were then removed, and mice were returned to their home cages. Once they returned to an alert state (15-20 min post isoflurane), mice were placed in the testing chamber directly from their home cage and video was recorded for 10 min. The rotating stage was gently rotated to enable videography of the face. Frames where the animal was facing away or during stage rotation were removed from videos using DaVinci Resolve software. All videos were analyzed from 0-6 min after removal of these segments. Rectal temperature was measured at 0.5-1 cm depth using the RightTemp Sensor rectal probe (Item # RT-0001-9, Kent Scientific), before infusion and after video capture.

### Video analysis

Our methods for behavioral analysis were adapted from previous work^11^. We used LabGym^24^, an AI-driven tool that automates behavioral tracking, to determine how activation of TRPV1 in the submandibular glands alters behavior. In particular, we measured the frequency and duration of 7 distinct behaviors: face swiping (paw lifting above the nose and then swiping at the face while the ear is pulled down), grooming (licking of front paws or body, scratching with hind leg), locomotion (movement within the chamber while all four paws remain on the floor), rearing (vertical positioning with paws on glass or outreached towards cylinder), resting (remaining in the same, relaxed position with ears upright and eyes open wide), sniffing (small, consistent snout-directed movements with the head moving snout-first either in the air or toward the bottom of the cylinder where it meets the floor), and grimace (orbital tightening, postural hunching, and ears pulled back). The LabGym Detector was trained to segment the mouse outline in all video recordings. The Detector was trained on 400 images from videos of mice in our behavioral acquisition setup. The images were annotated using Roboflow. Once the detector was trained to detect the mice in our videos, we generated behavior examples for training the LabGym Categorizer. The Categorizer was trained with 300-350 examples per behavior. Animation analyzer and pattern recognizer were set to level 5 and 2, respectively. Augmentations performed: horizontal flipping, random brightening, random rescaling. Wildtype mice that were given no experimental perturbation were recorded to obtain training examples of all behaviors except grimace. Grimace examples were obtained from videos of mice that were injected intraperitoneally with 125 mg/kg magnesium sulfate (CAS #7487-88-9, Millipore Sigma) dissolved in 0.9% sterile saline (CAS #7647-14-5, VWR) to induce abdominal constriction and pain-related behaviors^25^.The trained Detector and Categorizer were used to analyze our experimental videos. LabGym calculated the duration and frequency of each behavior and produced ethograms to visually illustrate behavioral differences between groups.

### Pilocarpine-stimulated saliva collection from submandibular glands

Mice were anesthetized via isoflurane administration (4% for induction and 2-3% for maintenance) using a SomnoSuite Low-Flow Anesthesia System (Kent Scientific) administered through a secured nose cone. Both Wharton’s ducts were cannulated, as described above, without an atropine injection. Cannulas were cut 1 cm protruding from the ducts just above the mandibular incisors. Mice were intraperitoneally injected with pilocarpine hydrochloride solution (Sigma-Aldrich, Catalog #P6503) (0.6 mg/kg) made in sterile saline to stimulate saliva production^8^. Saliva was collected only from Wharton’s ducts via the cannulas in an Eppendorf tube for 5 min once saliva secretion started. Volume was measured using a custom Hamilton syringe (30-gauge needle, Model #1702)

### Statistics

All statistical analyses were performed using GraphPad prism. Statistical methods, Mean or Median and Standard Error of the Mean (SEM) bars are described in the figure legends for individual experiments. Broadly, *t*-test (paired or unpaired) was used to compare groups. Statistical significance was defined as *P* values < 0.05.

## RESULTS

### Retrograde labeling captures trigeminal sensory neurons associated with the submandibular gland complex

We first sought to identify and define the trigeminal sensory neurons that are associated with the submandibular salivary gland. We reasoned that understanding the transcriptional classification would provide insight into the significance of this specialized sensory network. With this in mind, we introduced a cannula into the duct^19^ to deliver neuronal tracing agents to the submandibular gland complex. (**Fig. 1A**). Delivery of trypan blue verified successful infusion and gland targeting following experiments (**Supplementary Fig. 1**). For multiplex transcriptomic class mapping, we used ductal infusion of AAV6(F129L)/2-*hSyn1*-Cre to drive Cre-expression from a pan-neuronal *hSyn1* promoter and identified robust tdTomato^+^ labeling within a subgroup of sensory neurons of the ipsilateral ganglia (**Fig. 1B**). Neurons were distributed widely across the surface of the ganglion in both the rostro-caudal and mediolateral dimensions. To investigate the molecular identity of the retrogradely-labeled tdTomato^+^ trigeminal neurons, we applied multiplex *in situ* hybridization (ISH)-based classification^21,26^. Here, sections of trigeminal ganglia were evaluated using an 8-probe panel designed to distinguish major somatosensory classes (C1-C13 via probing for *Fxyd2, Calca, Trpa1, Tmem233, Trpm8, Mrgprd, S100b, and Nppb*) (**Fig. 1C**). Next, a machine learning algorithm was applied to predict the distinct transcriptomic profiles of tdTomato*^+^* cell bodies based on combinatorial labeling (**Fig. 1D**). Our analysis revealed that neurons innervating the submandibular gland’s complex primarily represented three distinct populations and defining genes: C4 Aβ mechanoreceptors, calcium-binding protein ß*, S100b* (33%, 159/473 neurons); C6 Aδ nociceptors, *S100b* and neuropeptide Calcitonin gene-related peptide, *Calca* (19%, 90/473 neurons); and a combination of C7-10 Aδ and C-type nociceptors and transient receptor potential vanilloid 1, *Trpv1* (31%, 143/473 neurons). While expression of *S100b* and *Calca* were confirmed as part of our 8-probe panel, we performed secondary validation for *Trpv1* expression. Here, ISH revealed that *Trpv1^+^* cell bodies comprised the majority of all labeled sensory TG neurons that innervated the submandibular gland’s complex (46%, 218/473 neurons, **Fig. 1E**). Overall, these data demonstrate distinct TG somatosensory innervation of the submandibular gland’s complex with substantial innervation by nociceptors, with the majority expressing *Trpv1*^+^.

**FIGURE 1.**
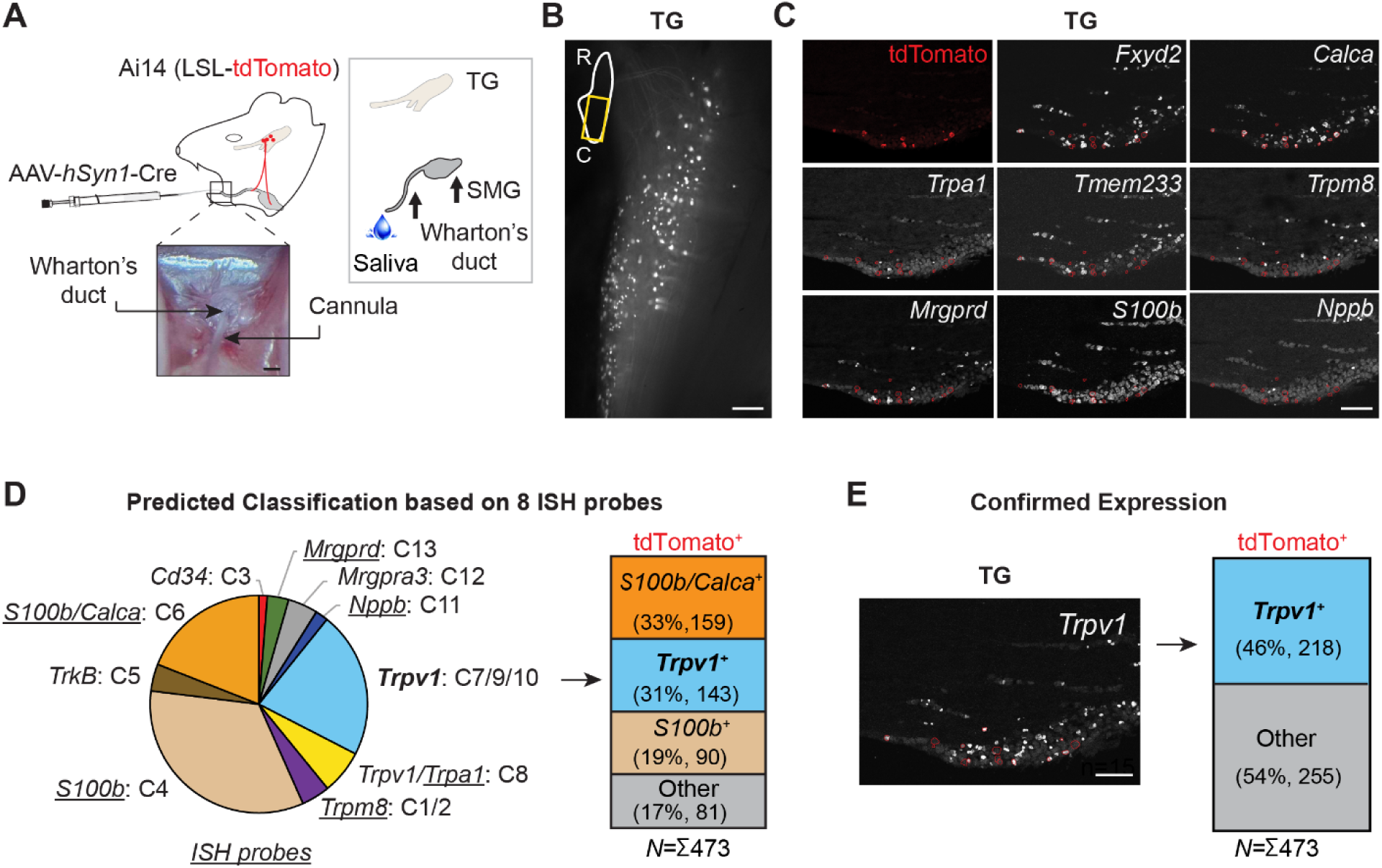
Retrograde labeling captures trigeminal sensory neurons associated with the submandibular gland complex. **(A)** Schematic illustrating placement of a cannula into the mouse submandibular gland’s Wharton’s duct and infusion of an AAV-*hSyn1*-Cre viral vector into an Ai14 Lox-Stop-Lox (LSL) tdTomato reporter mouse. Saliva flows out of the Wharton’s duct, which connects the SMG with the oral cavity. Scale bar, 200 µm. **(B)** Representative fluorescent (black and white pseudo colored tdTomato) image of dorsal TG from an AAV-*hSyn1*-Cre infused Ai14 mouse, as illustrated in (A). Scale bar, 200 µm. **(C)** ISH of 8 probes in sections of the tdTomato^+^ TG isolated in (B). Scale bar, 200 µm. **(D)** Pie chart reflecting number of TG neurons expressing tdTomato and ISH probes in each predicted neuronal class, as outlined in (C). A total of 473 neurons were counted across 3 (female and male) biological animals. Table depicts overall percentage and actual number of tdTomato-labeled TG neurons in the most abundant classes. **(E)** ISH for *Trpv1* on tdTomato-labeled TG neurons, as seen in (B), and quantified as an overall percentage of all tdTomato^+^ TG neurons. Scale bar, 200 µm. **Abbreviations:** TG, trigeminal ganglia. SMG, submandibular gland. R, rostral. C, caudal. ISH, *in situ* hybridization

### *Trpv1^+^* trigeminal sensory neurons innervate the Wharton’s duct

Given the robust, predominant, representation of *Trpv1*^+^ innervating TG neurons, and their well-studied role in deep tissue structures (e.g., cough, pain, mucus secretion)^27,28^, we focused our subsequent morphological and functional investigations on this population. To begin to distinguish the localization of *Trpv1*^+^ somatosensory fibers within the gland complex (i.e., gland body and Wharton’s duct), we used a genetic strategy (i.e., *Trpv1*-Cre;Ai140) to constitutively mark *Trpv1^+^* cells with GFP (**Fig. 2A**). Within the submandibular gland body, sparse GFP^+^ neuronal puncta were visible in the stroma surrounding the ECADHERIN^+^ epithelia (**Fig. 2B, Supplementary Fig. 2B**). However, this low level of fiber density seemed inconsistent with our observation of strong retrograde labeling of somata within the TG (**Fig. 1B**). We next turned our attention to the Wharton’s duct, which extends from the gland to provide final saliva transportation before its release into the oral cavity (**Fig. 1A**). In stark contrast with the gland, we observed a robust, concentrated, meshwork of GFP^+^ sensory terminals tracking prominently along the Wharton’s duct (**Fig. 2C**). Our genetic (*Trpv1*-Cre;Ai140) labeling approach also revealed the extent of local cellular expression of *Trpv1* in the epithelial compartment of the gland (**Fig. 2B**). However, GFP^+^ cells were sparse, often localized within a specific gland lobule (**Fig. 2B**, arrow), and only comprised 1.8 ± 1.2 % of all epithelial cells (**Fig. 2B**).

**FIGURE 2.**
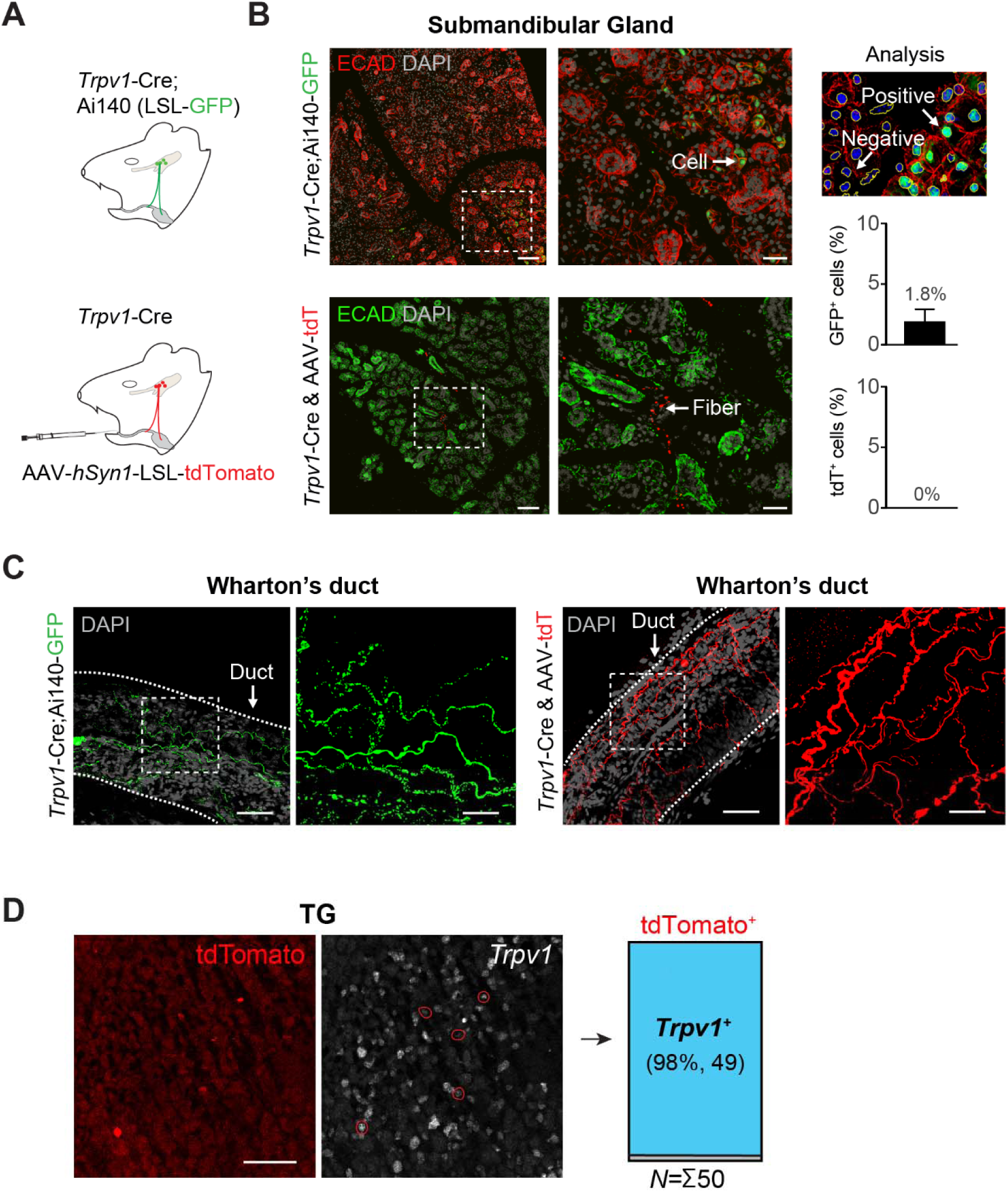
*Trpv1^+^*trigeminal sensory neurons innervate the Wharton’s duct. **(A)** Schematic of approaches used to label *Trpv1^+^* cells: (top) genetic crossing of *Trpv1*-Cre with Ai140 (LSL-GFP) mice, and (bottom) ductal infusion of AAV6.2-*hSyn1*-LSL-tdTomato into *Trpv1*-Cre mice. **(B)** Low and high-power confocal images of GFP and tdTomato expression in the submandibular glands, as depicted in (A) (*n* = 3 mice per condition). Tissue was counterstained with epithelial marker ECADHERIN (ECAD) and nuclei (DAPI). Scale bars, 100 and 20 µm. AI-mediated analysis of labeled cells. Graphs depict the percentage of GFP^+^ or tdTomato^+^ epithelial cells in the gland. **(C)** Confocal images of *Trpv1*-Cre mediated innervation of the Wharton’s duct (A). High power images represent insets of the dashed squares. Scale bars, 10 and 20 µm. (**D**) tdTomato^+^ neurons in the TG from *Trpv1*-Cre and AAV-*hSyn1*-LSL-tdTomato mice. Scale bar, 100 µm.

Because the constitutive *Trpv1*-Cre;Ai140 genetic mouse labeling not only marks *Trpv1*^+^ cells but also their prospective progeny since ontogenesis, we returned to AAV-mediated labeling to ensure that innervation represented *Trpv1*^+^ neurons at adulthood. Crucially, we found that AAV6.2-*hSyn1*-LSL-tdTomato infusion directly into the Wharton’s duct of adult *Trpv1*-Cre mice yielded tdTomato^+^ fibers with consistent gross anatomical distribution across the gland and Wharton’s duct’s architecture, mirroring the genetic labeling (*Trpv1*-Cre;Ai140) approach. Namely, few tdTomato^+^ fibers were seen within the submandibular gland (**Fig. 2B**), whereas a dense, mesh-like, innervation pattern prominently surrounded the Wharton’s duct (**Fig. 2C**). As expected, no tdTomato^+^ glandular epithelial cells were observed using our viral vector tracing (**Fig. 2B**), illustrating strong neuronal specificity of the *Trpv1*-Cre with AAV-*hSyn1* method. Postmortem *in situ* hybridization on corresponding trigeminal ganglia confirmed specificity of the AAV-mediated *Trpv1*^+^ neuronal fiber tracing: almost all of tdTomato^+^ somata expressed endogenous *Trpv1* transcript (98%, 49/50 neurons, **Fig. 2D**). Taken together, these findings demonstrate *Trpv1*^+^ trigeminal sensory innervation is enriched around the Wharton’s duct. Furthermore, these experiments demonstrate the neuronal specificity of our AAV-mediated tracing and support our use of a genetic reporter for evaluating *Trpv1*^+^ somatosensory neurons in downstream functional experiments.

### Ductal infusion of resiniferatoxin activates TRPV1^+^ trigeminal sensory afferents

The presence of *Trpv1*^+^ neuronal fibers in the submandibular gland’s complex suggests that these terminals may act as sensors in this environment. To test this and provide critical validation of TRPV1 function in these neurons, we performed an in vivo trigeminal calcium imaging technique (*Trpv1*-Cre;Ai95D)^11,22,29^ to visualize sensory activity during intraductal infusion of a TRPV1 agonist (**Fig. 3A**). Resiniferatoxin (RTX) is a potent vanilloid receptor agonist that induces sustained opening of the TRPV1 channel and neuronal activity^30^ and, thus, serves as a functional assay to determine the response of labeled *Trpv1*^+^ afferent neurons innervating the gland and Wharton’s duct. We found that ductal infusion of either vehicle (2 mM ascorbic acid, 0.25% Tween80 in ddH20) or RTX dissolved in vehicle evoked a temporal cascade of cellular activation within the trigeminal ganglion (**Fig. 3B**). These stereotyped recruitment of calcium responses could be reflective of fluid flowing into the Wharton’s and gland’s ductal architecture and locally encountering sequential receptive fields. RTX-induced responses showed an overall trend in increased number of responders (Mean, 19 vs. 40, **Fig. 3C**) with distinct response profiles. Whereas a subset of low-intensity responders (i.e., ΔF/F = 5-20) were present in both the vehicle and RTX group, RTX infusion evoked calcium responses with greater area under the curve (AUC, ΔF/F per min, Median = 643 vs. 1614, *P*<0.001) (**Fig. 3D, G**) and peak intensity (ΔF/F, Median = 15 vs. 24, *P*<0.0001) across all measured time (**Fig. 3E, G**). In addition, RTX infusion altered the functional relationship between peak intensity (ΔF/F) and AUC as determined by linear regression (**Fig. 3F**). Specifically, the slope for RTX-mediated responses was steeper than that of the vehicle, indicating a shift toward the recruitment of a different neuronal population with high intensity and activity (i.e., AUC) responses (**Fig. 3F**, circled dots are shown in G1-5). Overall, these data provide evidence that trigeminal afferents innervating the submandibular gland’s complex express functional TRPV1 ion channels that can be activated via ductal infusion and have the capacity to contribute to gland sensation and physiology.

**FIGURE 3.**
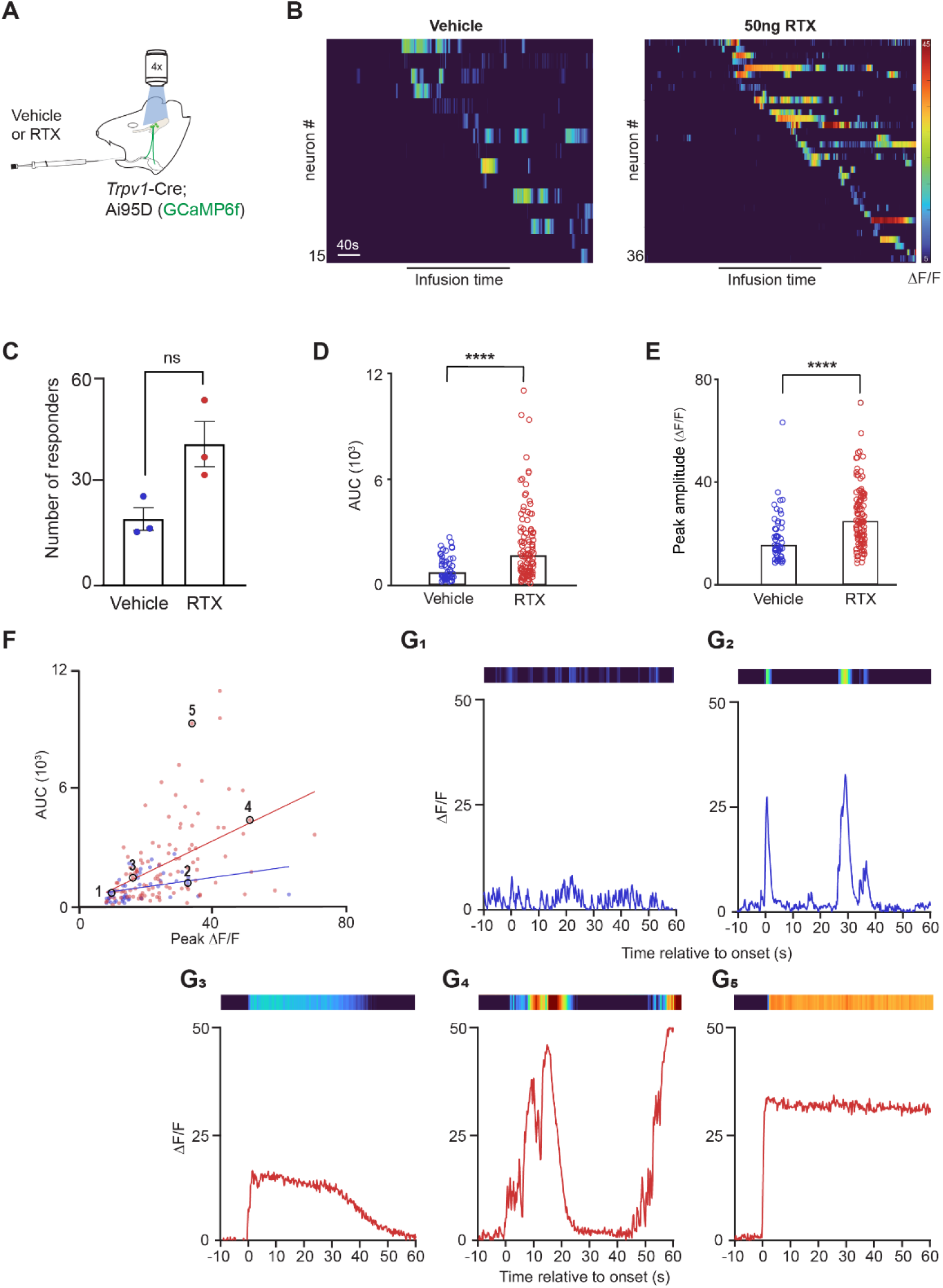
Functional activation of *Trpv1*^+^ trigeminal sensory neurons associated with the submandibular gland’s complex. **(A)** Experimental layout of a *Trpv1*-Cre;Ai95 (GCaMP6f) mouse being infused with a vehicle or RTX. Live imaging of the trigeminal ganglia measured GCaMP6f fluorescence as a readout for calcium fluxes. **(B)** Representative heatmap from single TG after vehicle (15 responding neurons) or RTX delivery (36 responding neurons). Vertical scale bars ΔF/F 5-40. **(C)** Number of responding neurons after vehicle or RTX delivery. Bar shows mean and error bars indicate the SEM. *n* = 6 mice, 3/condition, n.s*. P*>0.05, Mann-Whitney unpaired t-test. **(D)** Graph depicting the calcium response of vehicle and RTX responders as area under the curve (ΔF/F per minute), *P*<0.001, Mann-Whitney unpaired t-test. **(E)** Peak ΔF/F across the measured time of vehicle and RTX responders, *P*<0.001, Mann-Whitney unpaired t-test. **(F)** Linear regression graph of calcium response under the curve versus peak ΔF/F of vehicle and RTX responders, *n* = 173 neurons (54 vehicle, 119 RTX) from 6 mice (3 per condition). **(G)** 60 sec representative traces of single responders after vehicle or RTX delivery from black circles in (F). Vertical scale bars ΔF/F 0-50, corresponding heatmap above trace. **Abbreviations:** Resiniferatoxin, (RTX).

### Pharmacological activation of TRPV1 in the salivary gland complex drives orofacial pain

Salivary gland-associated pain is widely reported as a consequence of infection (i.e., sialadenitis^31^) or blockage from stones (i.e., sialolithisasis^32^). As our data revealed that *Trpv1^+^* afferents innervate the salivary gland and associated Wharton’s duct, and TRPV1^+^ neurons are generally involved with pain^33^, we analyzed whether TRPV1 could be a putative neuronal driver of gland complex pain. To test this directly, we activated TRPV1 channels via ductal infusion of RTX and evaluated subsequent evoked behaviors in freely moving mice (**Fig. 4A**). Because high global doses of TRPV1 agonists can engage central systemic thermoregulatory pathways^34^, we administered a low dose of RTX (2.5 ng) to avoid hypothermic effects after TRPV1 activation. To objectively evaluate and quantify mouse behaviors, the AI-driven analysis tool LabGym was leveraged^24^. Using independent control cohorts, the LabGym Categorizer was trained to classify baseline homeostatic behaviors (i.e., face swiping, grooming, locomotion, grimace, rearing, and sniffing) and establish an unbiased automated scoring system (**Fig. 4B**). In addition, definitive training examples of orofacial grimacing were generated from an independent wildtype cohort receiving intraperitoneal injections of magnesium sulfate, which reflects an established model of acute visceral pain^25,35^ (**Supplementary Fig. 2**). LabGym output generated frame-by-frame annotations, providing high-fidelity scoring of the animals’ behavior over time (**Fig. 4C**) (**Supplementary Video 1-2**). In our behavioral assay, 2.5 ng of intraductal RTX infusion drove a significant increase in grimacing phenotype well above the paired vehicle baseline (*P*<0.01, **Fig. 4C-D**). As expected at this low RTX dose, the grimacing phenotype was independent of any core body temperature reduction (**Fig. 4E**). Additionally, with increased RTX-evoked grimacing, there was a reduction in other typical control behaviors, most markedly grooming *(P*<0.01), locomotion *(P*<0.05), sniffing *(P*<0.05), and rearing *(P*<0.05) (**Fig. 4D**). Notably, vehicle-treated controls exhibited baseline fluctuations of grimacing behavior (**Fig. 4C**, red), indicating that even low volume intraductal fluid infusion is not entirely innocuous. Taken together, these data demonstrate that activation of TRPV1 channels through glandular infusion generates a general pain-related phenotype, suggesting that *Trpv1*^+^ trigeminal nociceptors may underlie some forms of pain within the submandibular gland complex.

**FIGURE 4.**
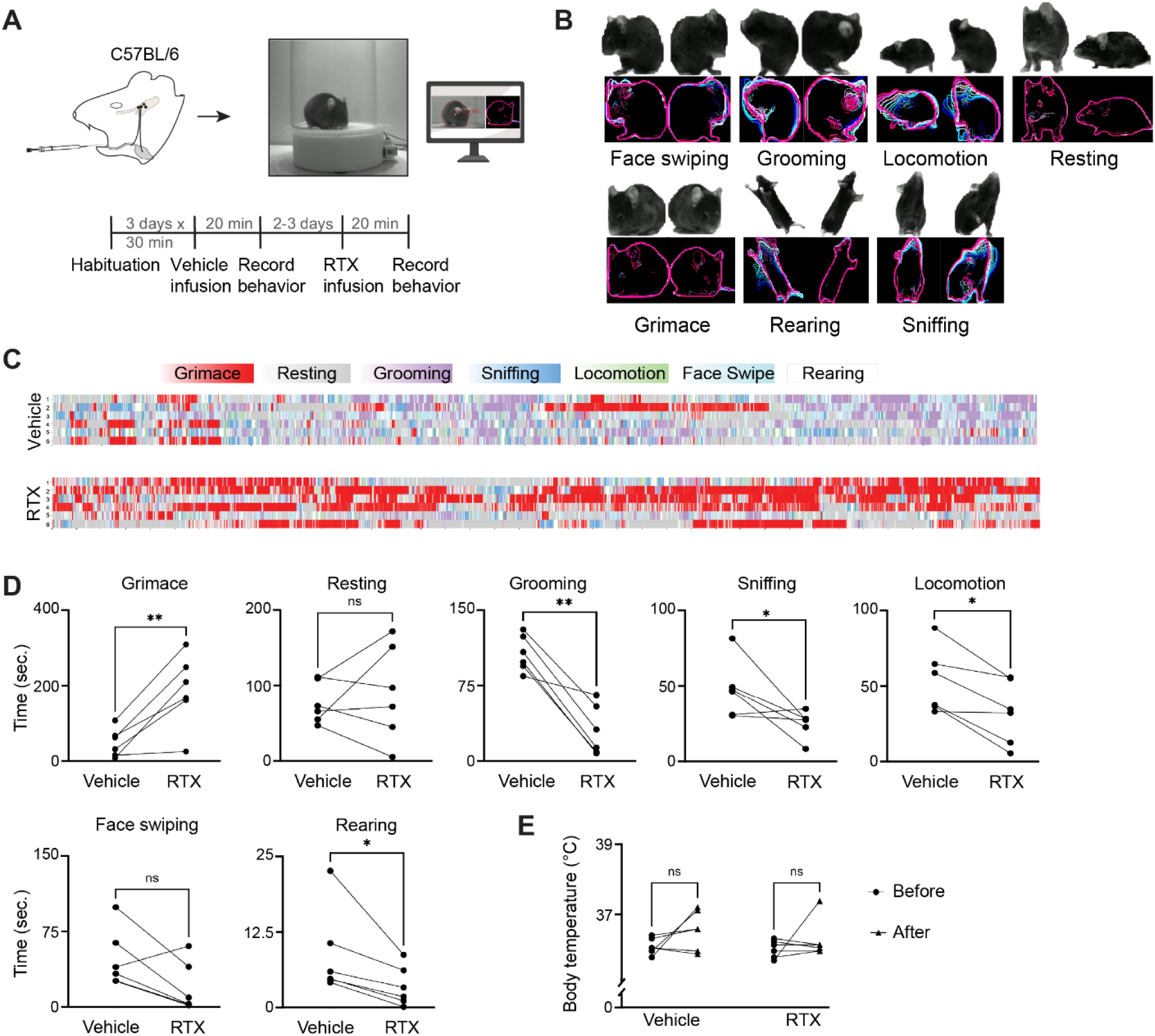
Pharmacological activation of TRPV1 in the salivary gland complex drives orofacial pain. **(A)** Schematic outlining the setup and timelines for infusion of vehicle and RTX into the salivary gland of wildtype mice for recorded behavioral experiments. **(B)** Representative examples showing video frame (top) and corresponding motion pattern images from LabGym (bottom) to illustrate animal behavioral categories. Overlaid color curves represent successive time points indicating behavior dynamics. **(C)** Behavioral recording (6 min) of wildtype animals treated with vehicle or RTX. Ethograms with each row representing an animal. Color bars indicate behavioral categories, with color reflecting the highest probability behavior. Time (*x* axis, s) and with each row (*y* axis) a single animal. *n* = 6 paired animals (3 male, 3 female). **(D)** Graph depicting the level of animal behavior between the vehicle-and RTX-treated animals from (C). Each connected dot represents the same animal pre-treated with the vehicle and then treated with RTX. Grimace (*P*<0.01), resting (n.s. *P*>0.05), grooming (*P*<0.01), sniffing (*P*<0.05), locomotion (*P*<0.05), face swiping (n.s. *P*>0.05), rearing (*P*<0.05), paired t-test. (**E**) Relationship between body temperature changes and level of grimacing in the same vehicle-and RTX-treated animal. **Abbreviations:** Resiniferatoxin, (RTX).

### Ablation of *Trpv1^+^* trigeminal sensory neurons disrupts secretion of saliva

As peripheral nociceptors are increasingly recognized as critical interoceptors that regulate aspects of organ physiology and homeostasis^4^, we investigated whether *Trpv1*^+^ afferents innervating the submandibular salivary gland and Wharton’s duct contribute to organ function. To test this, we ablated *Trpv1*^+^ cells using bilateral infusions of AAV-Diphtheria Toxin (DTA) or resiniferatoxin (RTX) into the Wharton’s ducts of *Trpv1*-Cre;Ai140 GFP reporter mice (**Fig. 5A**). Specifically, we introduced high dose resiniferatoxin (50 ng)^36^, which results in excitotoxicity and the rapid loss of local TRPV1-expressing terminals and cells. To complement this approach, in a separate cohort of mice, we administered AAV6.2-*hSyn1*-LSL-DTA to further restrict ablation to neurons actively expressing *Trpv1* (and Cre). Within the peripheral nervous system, expression of *Trpv1* is specific to somatosensory neurons^34,37^, thus this complementary AAV-DTA-based approach spares non-neuronal cells and specifically dissects the role of trigeminal afferents. Upon histological evaluation, the body of the submandibular glands of treated animals showed no overt alterations in morphology (**Fig. 5B**). Subsequent immunohistochemical evaluation of GFP-expressing fibers along Wharton’s duct (*Trpv1*-Cre; Ai140) confirmed that intraductal delivery of both RTX and AAV-DTA resulted in substantial local denervation compared to control mice (**Fig. 5C**). Changes were also observed in the Wharton’s duct (ECADHERIN^+^) epithelial architecture after *Trpv1*^+^ neuron ablation. The epithelial cells along the main duct exhibited altered morphology, suggesting disruption of epithelial organization and tissue architecture (**Fig. 5D**). AI-assisted image analysis of *Trpv1*^+^ fibers and innervation network density revealed significant differences between conditions. Both ablation strategies (DTA and RTX) resulted in a reduction in the amount of TRPV1^+^ fibers (DTA and RTX, *P*<0.01), and, consequently, a decrease in the innervation network density (*P*<0.01 and *P*<0.05, respectively) (**Fig. 5E**).

**FIGURE 5.**
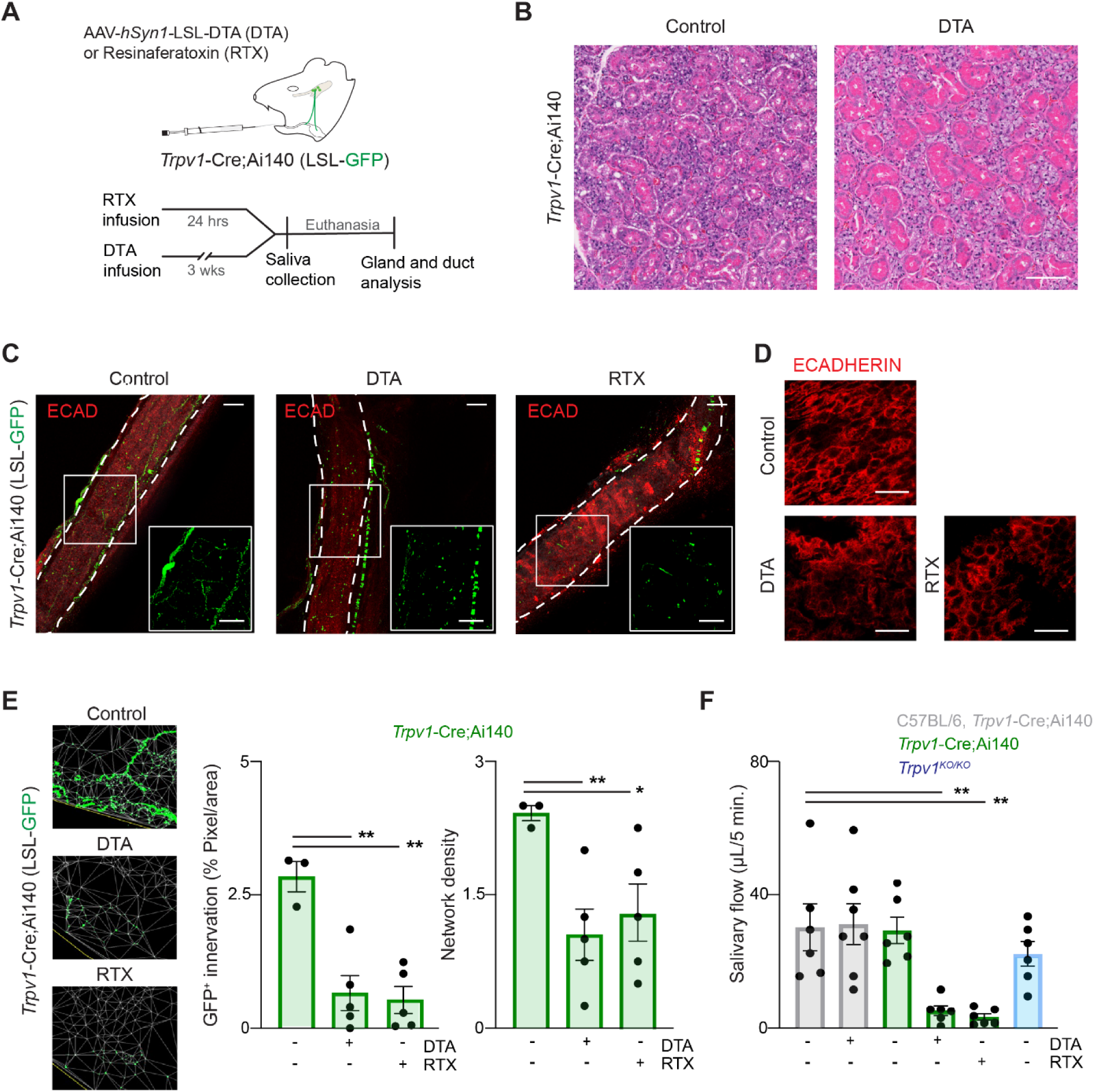
Ablation of *Trpv1^+^*trigeminal sensory neurons disrupts secretion of saliva. **(A)** Schematic outlining the experimental setup wherein RTX or AAV-*hSyn1*-LSL-DTA is infused in transgenic mice. **(B)** Histological hematoxylin-eosin images of a submandibular glands showing healthy states of acini in both control and DTA infused mice 3 weeks after infusion. Scale bar, 100 µm. **(C)** Confocal imaging and post-analysis of the presence of *Trpv1*^+^GFP^+^ neuronal fibers around the Wharton’s duct after AAV-*hSyn1*-DTA or RTX infusion. *n* = 13 mice (3 controls, 5 RTX, 5 DTA). Scale bar, 100 µm. **(D)** Architecture of Wharton’s duct epithelial cells stained for epithelial marker ECADHERIN after DTA or RTX infusion. Scale bar, 20 µm. **(E)** AI-assisted detection of innervation and generation of the innervation network mesh. Quantitative analysis revealed a significant reduction in the percentage of innervation in both DTA and RTX infused mice (both *P*<0.01), as well as a significant decrease in network density (DTA, *P*<0.01. RTX, *P*<0.05). Bars represent the mean, and error bars indicate the SEM. Unpaired t-test. **(F)** Saliva volume of various mouse strains and treatments with DTA or RTX. Bars show the mean, and error bars indicate the SEM. One way ANOVA, *n* = 6 mice ((3 male, 3 female)/condition), WT vs. DTA *P*<0.01, WT vs. RTX *P*<0.01.

Next, to test if a loss in *Trpv1*^+^ sensory innervation impacts the salivary gland complex’s ability to expel saliva, we collected cholinergic-stimulated saliva solely from the submandibular glands. With this technique, we reliably obtained saliva from control mice (Mean = 31 µL, *Trpv1*-Cre, Ai140, or C57BL/6). Saliva was not reduced by either infusion of AAV-DTA in the absence of Cre, ruling out nonspecific effects of viral vector ablation or strain variability. On the contrary, we found a profound reduction in total saliva volume obtained from both RTX and AAV-DTA ablation cohorts in both *Trpv1*-Cre;Ai140 males and females (<6µL, **Fig. 5F, Supplementary Fig. 3**). Interestingly, *Trpv1*-knock out (*Trpv1*-KO) mice did not reduce saliva flow, suggesting that disruptions in stimulated saliva secretion primarily stem from a loss in *Trpv1^+^* sensory innervation, but not TRPV1-mediated signaling (**Fig. 5F**). These combined data demonstrate that submandibular gland-innervating *Trpv1^+^* nociceptors are necessary for (stimulated) saliva secretion. Furthermore, our findings provide critical new insight into the diversity of nociceptor function, namely its contribution to essential exocrine gland physiology.

## DISCUSSION

Interoceptive somatosensory neurons have been increasingly identified in deep tissue structures throughout the body (e.g., lymph nodes^4^, kidneys^38^, and lungs^39^) where they contribute fundamentally to local tissue homeostasis and physiology. Earlier anatomical studies in rat^17,40^, sheep^41^, and dog^42^ provided preliminary insights that sensory afferents innervate salivary glands. In this study, we increase this body of work with compelling data to differentiate somatosensory innervation in the salivary glands’ complex. We provide in-depth anatomical and physiological analysis of *Trpv1*^+^ neurons that generate pain responses and serve as a crucial part of the gland’s stimulated secretory function.

Our *in situ* hybridization classification demonstrated that the mouse submandibular gland’s complex is innervated by a heterogenous population of trigeminal somatosensory neurons, of which *Trpv1*^+^ fibers predominate (31-46%). Apart from *Trpv1*, two other main classes were present in our findings: *S100b/Calca* (C6) and *S100b* (C4). Both classes represent large diameter neurons wherein C4 are mechanosensing^21^ (i.e., myelinated A-type low threshold fast-conducting mechanoreceptors that detect innocuous stimuli, such as touch) and C6 represent peptidergic Aδ nociceptors that rapidly detect pain^21^. Future work will investigate whether these neurons correspond to a possible mechanosensory role in sensation^18,43^.

One striking feature of the trigeminal *Trpv1^+^* afferents targeting the mouse submandibular gland’s complex is distinct anatomical compartmentalization. Earlier work that aimed to localize rat sensory afferents by CGRP and Substance P protein expression found CGRP^+^/Substance P^+^ nerves to be in close contact to ducts and blood vessels within the glands^44–46^. Using sophisticated genetic *Trpv1*-Cre mouse drivers with reporter mice or AAV-specific *hSyn1*-driven neuronal labeling, we also identified TG-derived *Trpv1^+^* afferent fibers localize in the submandibular glands’ stroma. However, they targeted the Wharton’s duct more robustly than the gland parenchyma, suggesting they may serve a larger role in this extended ductal environment.

Apart from *Trpv1^+^* fibers in the gland’s complex, our data also provide insight into the relative proportion of *Trpv1* in the epithelial parenchyma. We found this population was very scarce (1.8%) and was only locally present within specific lobules. These findings are in stark contrast with earlier immunostaining efforts for TRPV1 in rat parotid and sublingual glands^47^ and rat/human/rabbit^47–49^ submandibular glands where a large population of myoepithelial, acinar, and ductal cells appeared TRPV1^+^. Regardless, in our studies, neuronal TRPV1 specificity was facilitated using the *Trpv1*-Cre driver mouse and AAV-*hSyn1* model that allowed us to uniquely manipulate *Trpv1^+^* afferents without interference of any *Trpv1^+^* epithelia.

Our calcium imaging data provide key insights into the signaling stemming from trigeminal *Trpv1*-lineage derived sensory neurons innervating the glands’ complex. *Trpv1^+^* neurons associated with the gland’s complex demonstrated multiple distinct activity patterns upon RTX activation that correlated with prolonged, high intensity, calcium influxes. Our data further support that *Trpv1^+^* neurons act as one of the drivers of nociception within the gland’s complex system. Calcium responses to RTX further revealed that somatosensory fibers respond directly to chemical constituents infused into the ductal system. The lack of direct sensory innervation within the lumen indicates that the agonist RTX must permeate the epithelial barrier to activate fibers. The latter findings validate that ductal infusion can be used to modulate TRPV1 and other TG somatosensory neurons.

Our behavioral assays demonstrate for the first time that direct, peripheral, stimulation of the *Trpv1^+^* sensory population innervating the submandibular gland’s complex drives overt pain in mice. We found that noxious activation of the gland’s complex induces a robust grimacing phenotype. This finding further supports the general idea that pain stemming from various deep tissues can elicit conserved facial responses^25^. Pain is a common complaint in patients with sialolithiasis in the Wharton’s duct and smaller ducts of the gland^50^, illustrating both mechanoreceptive and nociceptive sensing can be activated. In particular, we were intrigued by our observations that the response to vehicle infusions was not only seen in calcium imaging but also in behavioral assays. These data suggest that infusion of any solution is not completely innocuous and may elicit a mechanical response due to changes in pressure. In addition, it cannot be ruled out that *Trpv1*^+^ innervation into the gland’s complex could also detect mechanical stimuli from the (vehicle or RTX) infusion. Altogether, our framework establishes an innovative platform for evaluating pain associated with the gland’s complex *in vivo*, providing an unbiased behavioral pipeline to discover and evaluate novel and targeted pain relief therapies.

In contrast to *Trpv1* activation, our ablation experiments (via DTA or high dose RTX) revealed an unexpected, fundamental, requirement for nociceptors in cholinergic-stimulated (i.e., pilocarpine) saliva secretion. Ablation of *Trpv1^+^* afferents significantly reduced stimulated saliva flow from the submandibular gland and was also associated with apparent changes in the Wharton’s duct global architecture. Our data also reveals that removal of TRPV1 (i.e., *Trpv1*-KO mouse) did not influence the submandibular glands’ stimulated saliva flow, indicating TRPV1 signaling is not directly correlated with saliva secretion. These results are in line with another study using *Trpv1*-KO mice wherein no change in whole saliva flow (from all salivary glands) was observed as compared with controls, even not after (i.p.) delivery of capsaicin^51^. Based on our findings, stimulated saliva flow depends on the structural presence of the sensory fibers rather than TRPV1-channel signaling itself. Furthermore, our findings suggest the presence of unidentified intermediate neuronal molecular factors that promote gland function. In other peripheral tissues, subsets of peptidergic *Trpv1^+^* nociceptors are known to exhibit baseline firing and secrete neuropeptides to modulate epithelial proliferation, vascular tone, and tissue homeostasis^52^. We propose that these duct-associated *Trpv1^+^* neurons might maintain a baseline level of low-threshold activity that drives the continuous release of trophic factors necessary to preserve the epithelial cell-cell interface. Future studies will be needed to uncover the specific transcriptomic and/or secretory repertoire of these duct-innervating *Trpv1^+^* fibers. Furthermore, understanding the downstream signaling cascade within surrounding non-neuronal cells will shed light on the molecular mechanisms that preserve exocrine gland physiology and may be leveraged for tissue regeneration.

In conclusion, our study indicates that *Trpv1^+^* neurons associated with the salivary glands’ complex respond to acute ductal-associated fluid infusions, drive pain behaviors, and facilitate saliva secretion into the oral cavity. Intraductal activation of these trigeminal afferents represents a model for pain in the gland complex that can be leveraged to develop targeted analgesics. Further experiments will examine how steady-state sensory activity can be uncoupled from acute nociceptive signaling to preferentially drive (or inhibit) these independent functional outputs. Overall, our investigation indicates that duct-associated *Trpv1^+^* neurons in the submandibular salivary glands represent a distinct subclass of deep-tissue afferents that are uniquely tuned to execute dual physiological roles, serving as both an acute sensor of pain and an indispensable mediator of daily organ homeostasis and function.

## Supporting information

Video - Example LabGym-annotated videography demonstrating wildtype mouse with vehicle infusion

Video - Example LabGym-annotated videography demonstrating wildtype mouse with RTX infusion

## ACKNOWLEDGEMENTS

This work is supported by NIH grants R01 DE032345 (J.J.E. and I.M.A.L.), F31 DE034282 (D.N.C), and T32 DC000011 (D.N.C). We thank Samuel Zahner for computational technical assistance in designing and revising custom code; Alvin Chiu for initial insight on calcium imaging analysis; Dr. Audrey Seasholtz, Dr. Bo Duan, and Dr. Ada Eban-Rothschild for valuable suggestions; and members of the Emrick lab for continued support and feedback.

## AUTHOR CONTRIBUTIONS

All authors gave final approval and agreed to be accountable for all aspects of the work. J.J.E., I.M.A.L., and D.N.C. conceptualized the study. D.N.C. performed all AAV cannulation and Ca2+ imaging experiments with initial support with analysis from A.R.G. R.E.W. and D.C. performed cannulation experiments for behavioral experiments, R.E.W. and K.M.M. obtained behavioral recordings and conducted analysis with D.N.C. I.A.V. contributed to initial experiments of gland isolations. J.A. isolated salivary ducts and glands and performed immunostaining with A.S. D.N.C. performed saliva ductal collection. D.N.C., and J.J.E. drafted the manuscript with critical edits from J.A. and I.M.A.L. The manuscript was finalized with input from all other authors.

## DECLARATION OF INTERESTS

Authors declare no competing interests.

## Notes

### Competing Interest Statement

The authors have declared no competing interest.

