## Supplementary figures and figure legends for "*Trpv1^+^*sensory innervation of the salivary gland drives pain and supports saliva secretion"

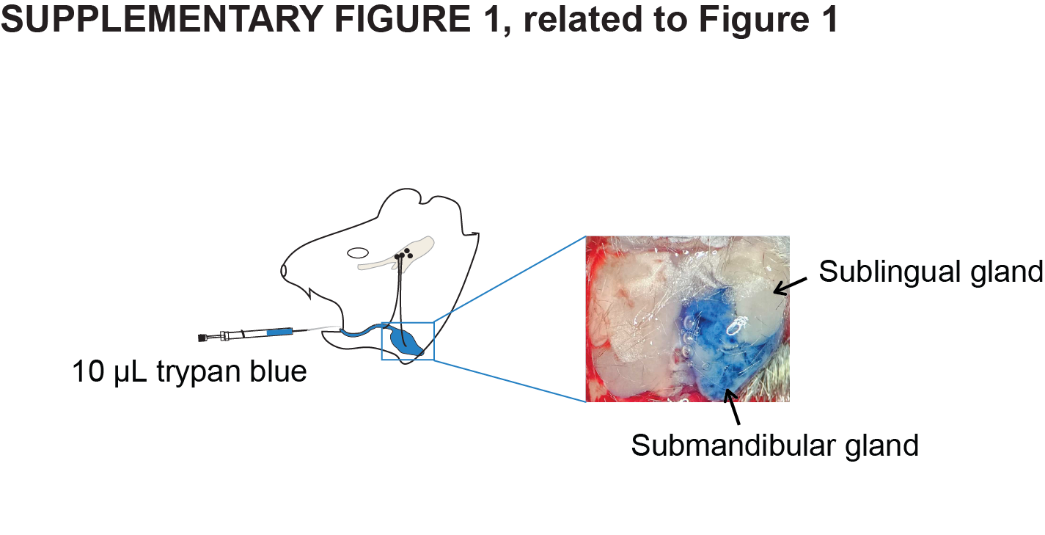

**SUPPLEMENTARY FIGURE 1, related to Figure 1**. **Retrograde ductal infusion.** Schematic of Trypan blue infusion into Wharton’s duct and SMG. Inset represents visual validation of localized dye delivery restricted to ipsilateral gland following cannulation and infusion.

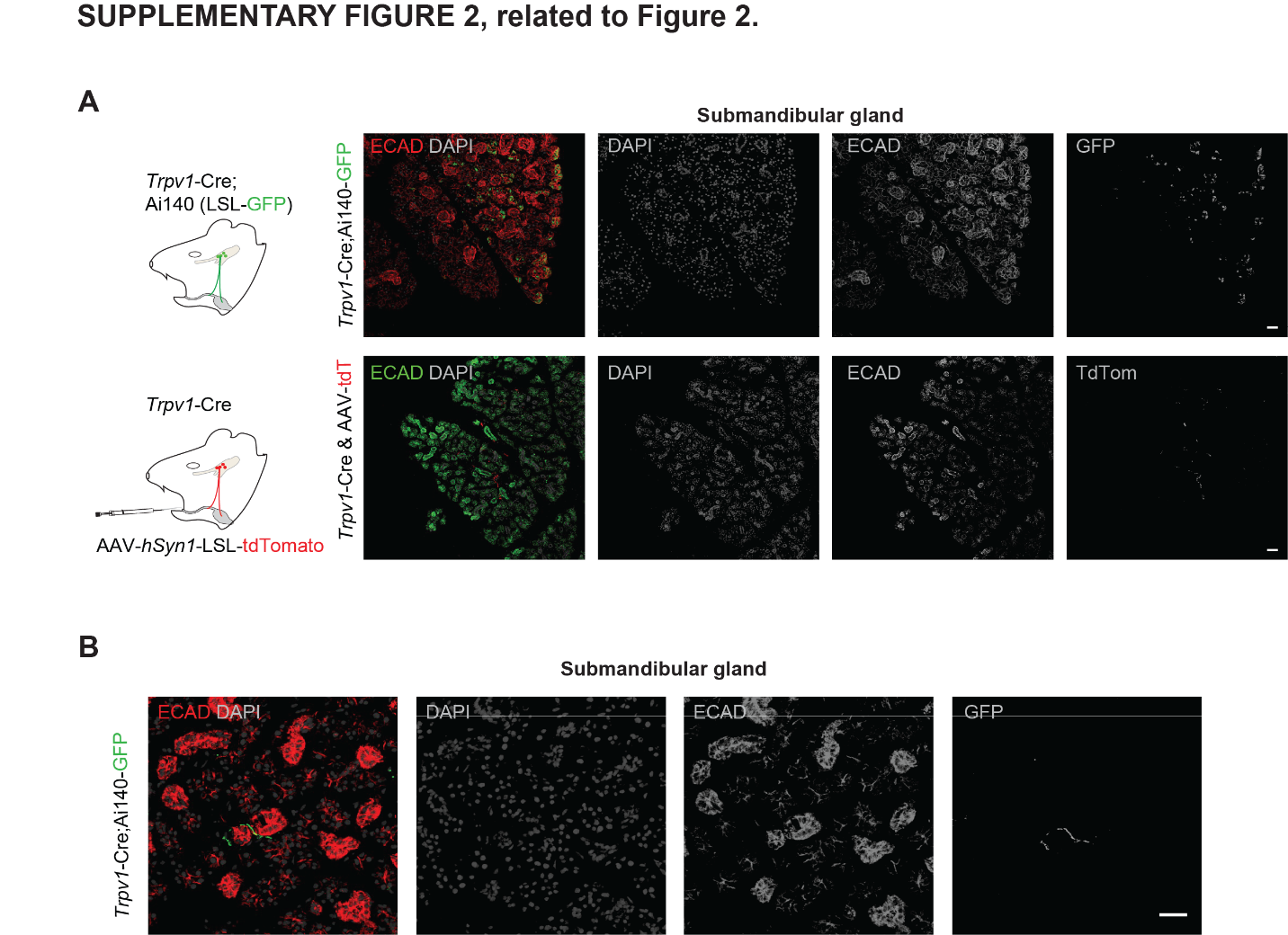

**SUPPLEMENTARY FIGURE 2, related to Figure 2**. ***Trpv1*-GFP^+^ cells in the submandibular gland.** **(A)** Images from Figure 2B illustrated as black and white images for each channel. **(B)** GFP^+^ fiber terminals in the submandibular gland of *Trpv1*-Cre;Ai140 mice.

**SUPPLEMENTARY VIDEOS 1-2. Video showing behavior of mice treated with vehicle or RTX.**

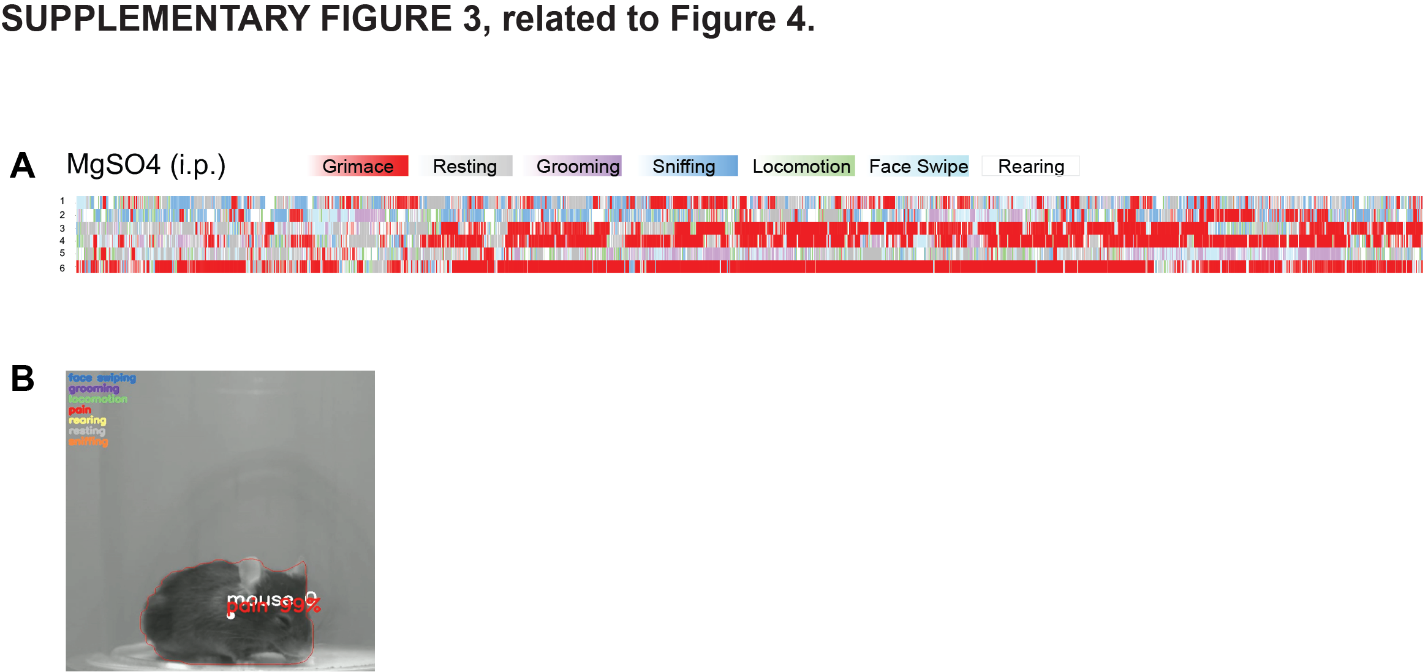

**SUPPLEMENTARY FIGURE 3, related to Figure 4**. M**agnesium sulfate pain model.** **(A)** Behavioral recording (6 min) of wildtype animals treated with i.p. injections of magnesium sulfate. Rastermap with each line representing an animal. Color bars indicate behavioral categories, with color intensity reflecting the probability of each behavior. Time (*x* axis, s) and with each row (*y* axis) a single animal. *n* = 6 (3 male, 3 female). **(B)** Annotated video frame from LabGym depicting confidence in categorizing pain expression.

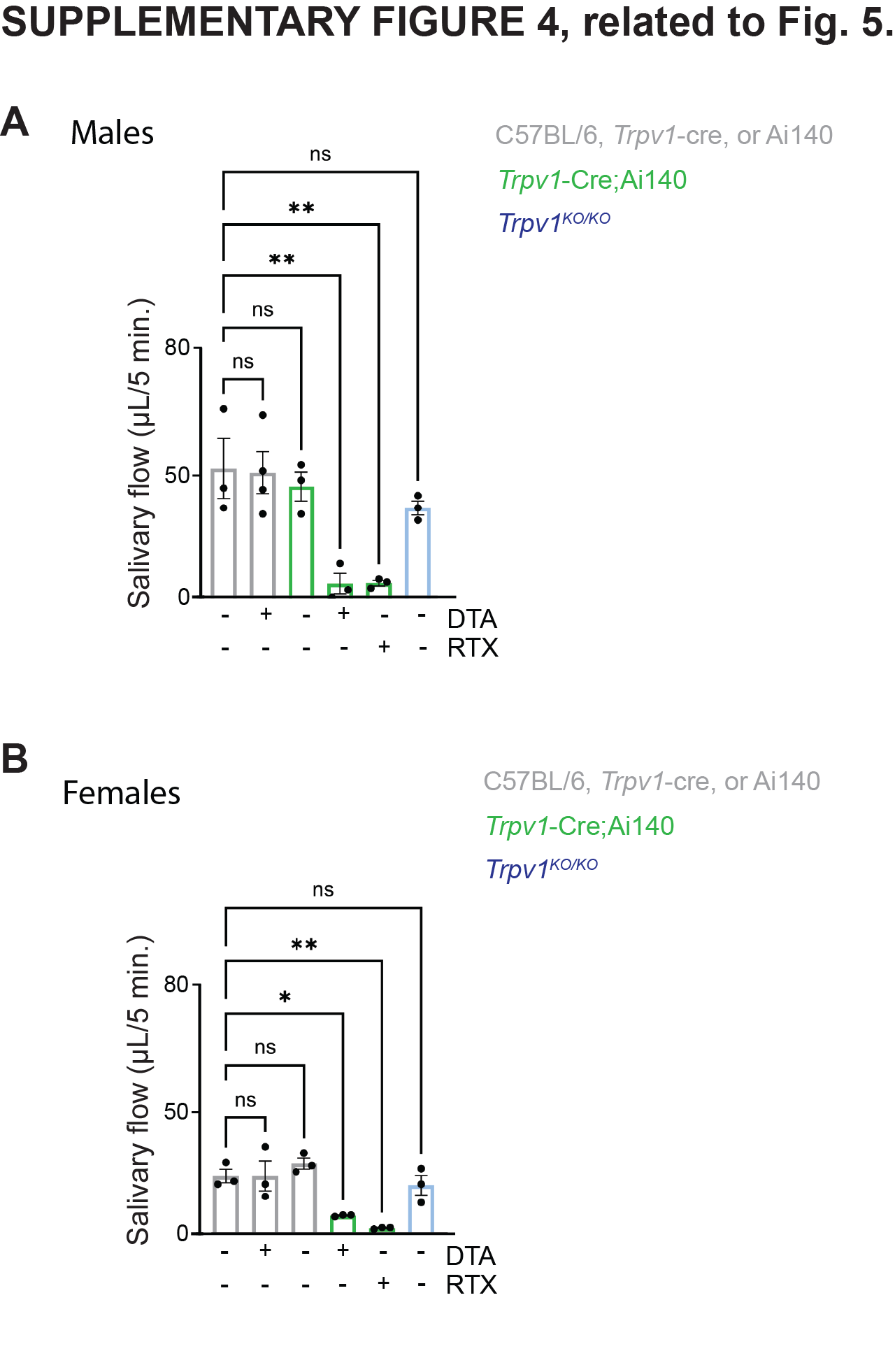

**SUPPLEMENTARY FIGURE 4, related to Figure 5**. **Salivary volume obtained following *Trpv1^+^* ablation.** Differences in saliva volume separated between male and female mice following DTA or RTX, as seen in Figure 5G. **(A)** Males, *n* = 3 mice/condition, WT vs. DTA *P*<0.01, WT vs. RTX *P*<0.01. **(B)** Females, *n* = 3 mice/condition, WT vs. DTA *P*<0.05, WT vs. RTX *P*<0.01. One way ANOVA.
